# The dynamics of SNARE complex assembly and disassembly in response to Ca^2+^ current activation in live chromaffin cells

**DOI:** 10.1101/2025.01.24.634783

**Authors:** Qinghua Fang, Ying Zhao, Dong An, Manfred Lindau

## Abstract

A SNAP25 based FRET construct named SCORE (SNARE COmplex REporter) has revealed a transient FRET increase that specifically occurred at fusion sites preceding fusion events by tens of milliseconds and presumably reflects vesicle priming. The FRET increase lasts for a few seconds until it is reversed. In those experiments, the FRET increase was found to be localized to areas <0.5 µm^2^ at sites of transmitter release as detected amperometrically using electrochemical detector arrays. Due to the localization to such small areas, it was unknown if the reversal of the FRET increase is due to local dispersion of high-FRET SCORE copies leaving the site after fusion and exchange with surrounding low-FRET copies, or if it reflects disassembly of the high-FRET complexes. To resolve this question, we performed whole-cell patch clamp pulse stimulation experiments, imaging the entire footprint of the cells in Total Internal Reflection Fluorescence (TIRF) excitation mode such that diffusional exchange between high-FRET and low-FRET copies does not produce a net FRET change. We show here that pulse stimulation of calcium currents results in FRET ratio transients with a time course very similar that related to fusion events. By comparing the kinetics of the FRET ratio decay with analytical and numerical diffusion simulation results, we show that the experimentally observed kinetics cannot be explained by diffusional exchange and conclude that the SCORE FRET ratio transients reflect incorporation of SCORE in SNARE complexes followed by SNARE complex disassembly. Experiments using Synaptobrevin 2/Cellubrevin double knock-out mouse embryonal chromaffin cells showed no pulse induced FRET change, indicating that the vSNARE is required for the incorporation of SCORE (or SNAP25 in wild type cells) in the SNARE complex during priming.

**Statement of Significance:** In chromaffin cells, SNAP25-based FRET constructs (SCORE) revealed transient FRET increases within <0.5 µm^2^ areas, preceding individual fusion events that reversed within seconds. It remained unknown whether this reversal stems from high-FRET complex disassembly or diffusion-mediated exchange with low-FRET complexes. Here, we performed whole-cell patch-clamp pulse stimulation with TIRF microscopy, imaging large ∼30 µm^2^ areas of the cell footprint. Calcium currents induced FRET transients with the same decay time constant of ∼1.5 s, significantly shorter than the time scale of diffusion. The SCORE FRET ratio thus reports in real time the dynamics of SNRE complex assembly and disassembly in live cells. Using Synaptobrevin 2/Cellubrevin double knock-out mouse chromaffin cells we show that vSNAREs are required for SNAP25 incorporation into SNARE complexes during priming.

## Introduction

The neuronal SNARE (Soluble NSF Attachment REceptor) complex is composed of the proteins Synaptobrevin 2 (Syb2 also known as VAMP2), Syntaxin 1 (Stx1), and SNAP25. These core complexes represent the major components of the fusion nanomachine and conformational changes in one or more SNARE complexes are thought to mediate the sequence of steps, from vesicle-plasma membrane docking and priming to the formation and dilation of fusion pores that allow for rapid transmitter release (1-4). Munc13 switches Stx1 to the open conformation and thereby recruits SNAP25 to the docking site preparing the vesicle for fusion in a Ca^2+^ dependent priming step (5) such that the vesicle can be released rapidly in response to stimulation, i.e. Ca^2+^ entry via voltage-gated Ca^2+^ channels. The development of a SNAP25 based FRET construct named SCORE (Snare COmplex REporter) incorporating CFP as FRET donor and Venus as FRET acceptor (6) or a variant using mCerulean3 as FRET donor named SCORE2 (7) at the N termini of the two SNAP25 SNARE motifs SN1 and SN2, respectively (Fig. 1A), has made it possible to monitor SNARE complex dynamics in live cells. We previously found that in chromaffin cells, under conditions of steady state priming, fusion, and release, a FRET increase precedes individual fusion events by ∼90 ms in bovine chromaffin cells (8) and by ∼65 ms in embryonal mouse SNAP25 knock-out cells (7). SCORE2 fully rescues fusion and properties of quantal release events in the SNAP25 knock-out cells, indicating that it is a fully functional replacement for SNAP25 (7). These FRET changes of the constructs named SNARE COmplex REporter (SCORE) and SCORE2, respectively, presumably indicate the incorporation of SCORE (or SNAP25 in wild type cells) in SNARE complexes during the priming step (Fig. 1B). Vesicle priming is associated with a transition to a tightly docked state with a vesicle-plasma membrane distance of 4-5 nm (9-13). It has been suggested that the transition to this tightly docked state requires synaptobrevin, syntaxin, and specifically SNAP25 (14) (Fig. 1C). The FRET increase reported by SCORE or SCORE2 is transient, reversing within a few seconds. Since the FRET increase was highly localized to a very small area <0.5 µm^2^, it was unknown if the reversal of the FRET increase was due to local dispersion of high-FRET SCORE copies leaving the site after fusion and exchange with surrounding low-FRET copies, or if it reflects disassembly of the high-FRET complexes. To avoid this ambiguity that comes from the analysis of localized FRET changes we designed experiments where a large area of the membrane is monitored such that diffusional exchange between high-FRET and low-FRET copies will not produce a net FRET change.

**Fig. 1.**
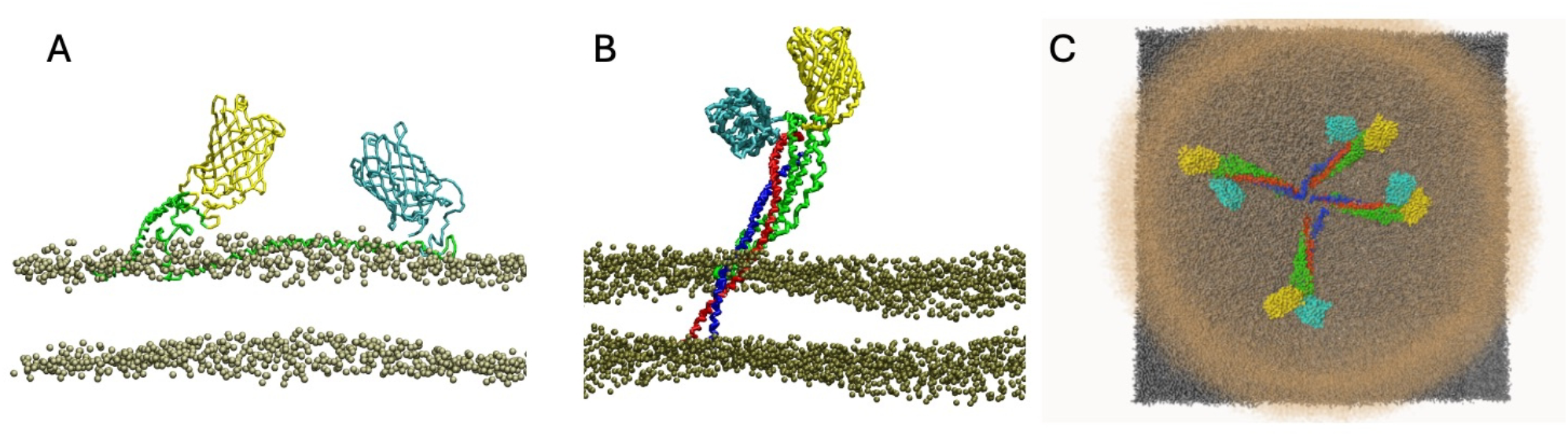
Illustration of the approach using SNAP25 (green) based FRET constructs incorporating CFP or mCerulean3 (cyan) as a FRET donor and Venus (yellow) as FRET acceptor at the N termini of the two SNAP25 SNARE motifs SN1 and SN2, respectively. The head groups of the lipids of plasma membrane are colored brown. SNAP25 alone is largely unstructured, and FRET efficiency is low (A). Upon incorporation into a SNARE complex (Syb2 and Stx1 colored blue and red respectively) it becomes highly structured and FRET efficiency increases (B). The top view of the proposed fusion system with a possible arrangement of multiple SNARE complexes incorporating SCORE in the tightly docked, primed state. Illustrations adapted from (7) (C). The synaptic vesicle and the plasma membrane are colored brown and black respectively.

It is well established that vesicle priming is a calcium dependent step (15). We therefore performed experiments stimulating calcium influx in chromaffin cells expressing SCORE or SCORE2 using pulse depolarization in whole-cell patch clamp experiments and monitored the FRET changes in a large ∼30 µm^2^ area of the cell’s footprint at the center of the cell in TIRF excitation mode. The experiments reveal that pulse stimulated calcium currents induce a transient FRET change in the large membrane area with a time course that is very similar to that recorded in the small (<0.5 µm^2^) areas preceding individual fusion events under steady state conditions. In particular, the exponential decay of the FRET ratio following the plateau of the FRET increase occurs with a virtually identical time constant indicating that the reversal of the FRET increase reflects the disassembly of the SNARE complexes that incorporated SCORE during priming. The results also reveal that the priming step indicated by the SCORE2 FRET change requires the presence of vSNAREs.

## Results

### Activation of calcium currents induces a transient FRET increase indicating a structural change of SNAP25

Since vesicle priming is a Ca^2+^ dependent process, we tested the hypothesis that Ca^2+^ influx via voltage gated Ca^2+^ channels stimulated by depolarization pulses should induce a FRET change in SCORE overexpressing cells. Bovine chromaffin cells expressing SCORE were voltage clamped in the whole-cell patch clamp configuration while membrane capacitance was recorded using the Lindau-Neher technique (16) and SCORE was excited using a 436/10 nm excitation filter in Total Internal Reflection Fluorescence (TIRF) excitation mode. Fluorescence emission from an ∼30 µm^2^ area of the footprint of the cell was imaged simultaneously in the CFP (465-495 nm) and Venus (520-550 nm) bands. Cells were stimulated with 25 ms pulses from −70 mV to +10 mV and sections starting 10 s before the pulse and ending 10 s after the pulse were extracted for analysis. Ca^2+^ currents in chromaffin cells show cell-to-cell variability, which created the opportunity to split the cells in two groups based on Ca^2+^ current amplitude into either a high Ca^2+^ current group “H” having Ca^2+^ current >25 pA (2625 pulses during 312 recordings from 252 cells) or a low Ca^2+^ current “L” group having Ca^2+^ current <25 pA (1421 pulses during 162 recordings from 127 cells). Within each group the traces were averaged (Fig. 2). In group H the averaged Ca^2+^ current in the time interval 5 to 25 ms was −115 pA (2. 1Ai). It induced a persistent average capacitance increase by 4.5 fF (Fig. 2A, Aii). In group L the average Ca^2+^ current (Fig.2 Bi) was undetectable, and the induced persistent capacitance increase was <1 fF (Fig. 2B, Bii).

**Fig. 2.**
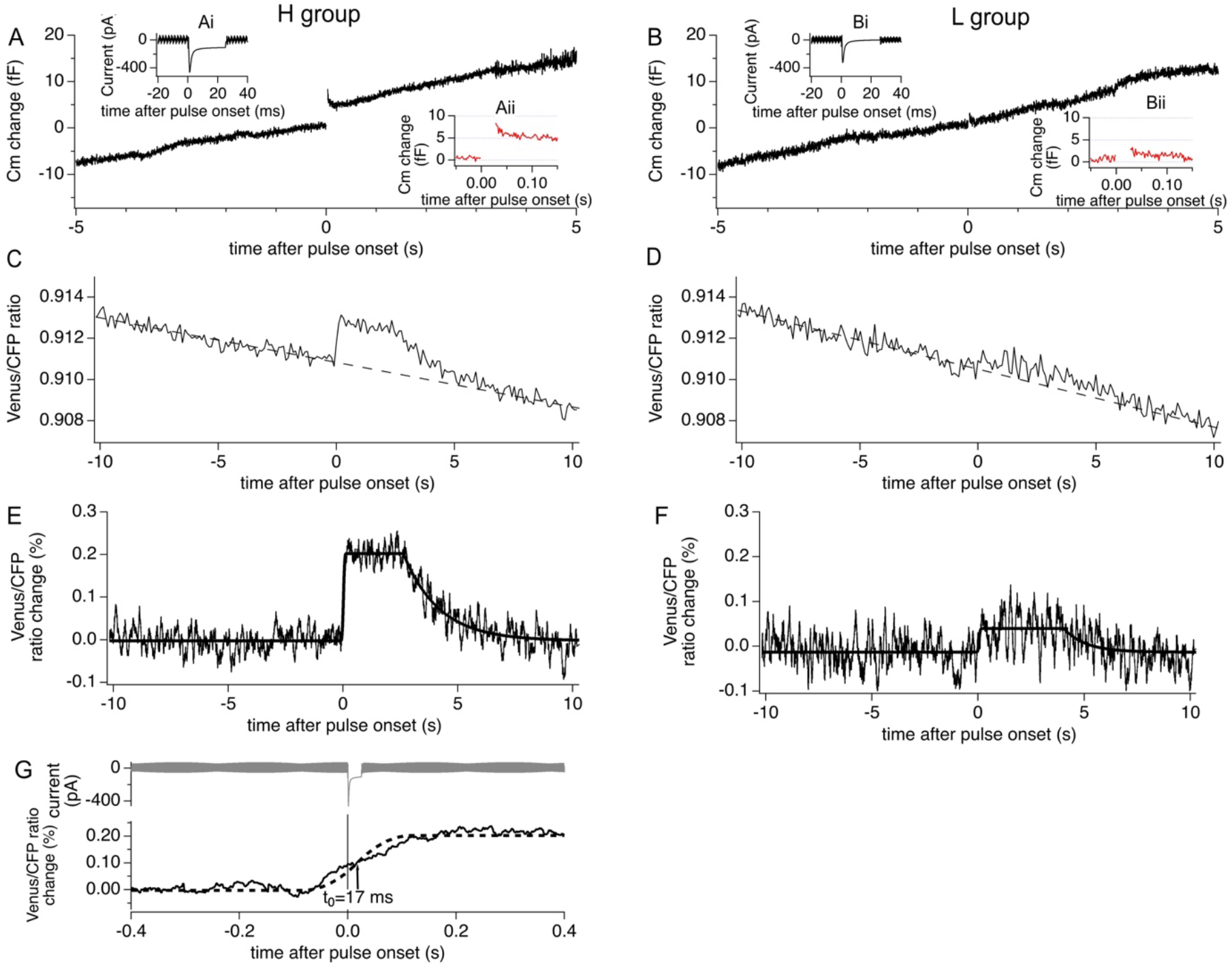
Stimulation of Ca^2+^ currents induces SCORE FRET changes in bovine chromaffin cells. Stimulations producing high Ca^2+^ currents (H group, left panels) (Ai) induce capacitance increase (A) also shown on expanded scale (Aii). Stimulations producing low Ca^2+^ current (L group, right panels) (Bi) induce no capacitance change (B, Bii). H group cells show transient FRET ratio increase (C), which is strongly diminished in L group cells (D). The FRET time superresolution traces after baseline subtraction (E, F) were fitted with the ECOM step response function modified to account for the decrease following the plateau by adding an exponential decrease back to baseline starting the end of the plateau. The ECOM filtered time course of the initial step response (G) indicates a time of FRET increase 17 ms after pulse start time. All traces are averaged traces from the respective groups.

The FRET ratio was calculated frame by frame as the fluorescence intensity ratio in the Venus/CFP bands. A clear transient increase in the FRET ratio is evident in the H group (Fig. 2C) but is profoundly reduced in the L group (Fig. 2D). For a detailed analysis of the FRET change kinetics, the FRET ratio frames were temporally aligned with 1 ms sub-frame time resolution to the stimulation pulse start time, which occurred randomly relative to the imaging frames. After the alignment, the intensity ratio traces were averaged using the Event Correlation Microscopy (ECOM) method, which provides a time super resolution trace of the average time course of the FRET ratio change low pass filtered with the ECOM step response function (17). This trace, after subtraction of the FRET ratio baseline drift (Fig. 2C, dashed line), shows a rapid increase followed by a plateau and a subsequent exponential decrease in the H group (Fig. 2E) and a much smaller transient in the L group (Fig. 2F). A fit with the ECOM step response function (Equation 9 in ref (17)) provides the amplitude of the FRET ratio change (Δ*R*_*max*_) and the time *t*_*0*_ relative to the time of pulse onset. To account for the decrease following the plateau, an exponential decrease back to baseline was added to the fit function, providing the time for the end of the plateau, *t*_*plateau*_, where the exponential decay starts and the time constant of the exponential decay *τ*_*decay*_. The fit results reveal that the average amplitude of the FRET ratio change is 0.205 ± 0.006% in the H group and 0.053 ± 0.006% in the L group and is thus ∼4 times smaller in the L group. The FRET ratio change is therefore clearly related to the size of Ca^2+^ influx generated by the depolarization pulse. The parameter *t*_*0*_ in the H group (Fig. 2G) is 17 ms after pulse start and indicates that the FRET ratio change occurs during the stimulation pulse.

### The SNAP25 conformational change indicated by the FRET change depends on the presence of vSNAREs

The ability to stimulate the formation of the primed state, reflected in the incorporation of SNAP25, or SCORE in this case, with Ca^2+^ current pulses makes it possible to study the role of the vSNAREs in this process even under conditions where fusion does not occur. Since both, Syb2 and Cellubrevin (Ceb) support vesicle fusion in chromaffin cells (18), we compared the results from Syb2/Ceb double knock-out embryonal mouse chromaffin cells (from here called DKO cells) with control wild-type embryonal mouse chromaffin cells (WT cells). In these cells we expressed SCORE2, which incorporates mCerulean3 as FRET acceptor instead of CFP in the original SCORE.

These experiments were performed to determine the relative size of the Venus/mCerulean3 FRET ratio changes in Syb2/Cellubrevin DKO compared to WT mouse embryonal chromaffin cells. For a robust response, cells were stimulated with 200 ms pulses from −70 mV to +10 mV (Fig. 3) and the SCORE2 FRET ratio was monitored in the cell’s footprint. Typically, only the first pulse and in some cases the second pulse generated a significant capacitance change in WT cells. Therefore, for the comparison with DKO cells, only the first pulse applied in each cell was used in the analysis. The average Ca^2+^ currents were unchanged in DKO cells compared to WT cells, as expected (Fig. 3A, B). The average capacitance increase stimulated by the pulse in WT cells was virtually absent in DKO cells (Fig. 3C, D). The average basal SCORE2 FRET ratio was averaged over 1.5 s preceding the pulse onset and the SCORE2 FRET ratio after the pulse was averaged over a 1.5 s time interval starting 0.3 s after the pulse onset. The basal FRET ratio was slightly reduced in DKO cells compared to WT cells (Fig. 3E, F) but this difference was not significant (p=0.38). The pulse induced increase in FRET ratio (Fig. 3E, G) in WT cells (0.0049 ± 0.0008, 10 cells) was absent in DKO cells (−0.00031 ± 0.00065, 13 cells). While it has previously been shown that vSNAREs are required for tight docking and priming in the synapse (12), this has not been the case in chromaffin cells. It was previously reported that docking of mouse embryonal chromaffin granules was unchanged in DKO cells (18) but in this analysis potential changes in tight docking (<20 nm distance) could not be examined. The results shown in Fig. 3E, G clearly demonstrate that the SCORE FRET changes indicating vesicle priming depend on the presence of vSNAREs.

**Fig. 3.**
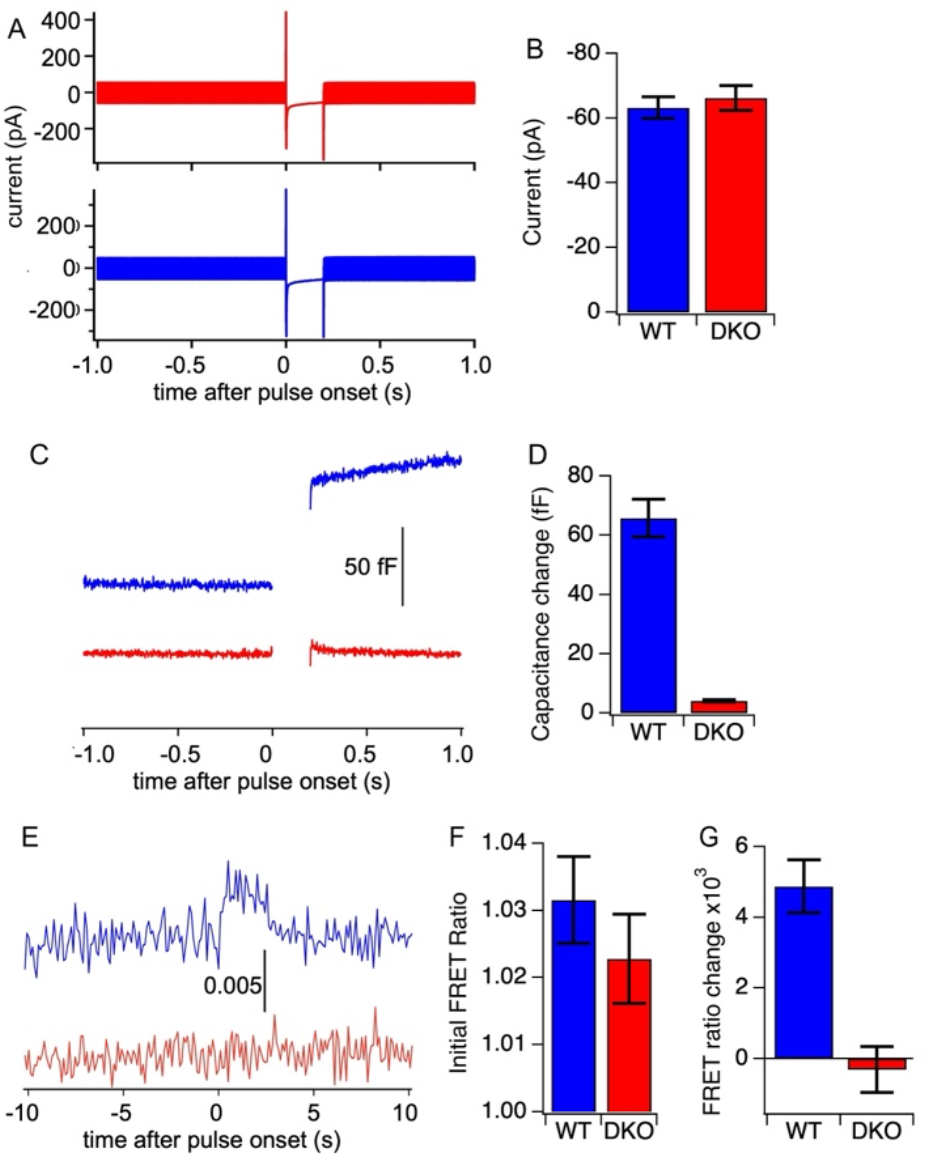
Syb2/Cellubrevin DKO cells show no pulse induced FRET ratio transient (DKO: n=13, WT: n=10). (A) Averaged Ca^2+^ currents in DKO and WT cells, (B) statistical analysis (WT: 63.2 ± 3.3 pA (mean ± SEM), DKO: 66.2 ± 3.9 pA, p=0.59). (C) Averaged capacitance traces from DKO and WT cells, (D) statistical analysis of capacitance changes (WT: 65.8 ± 6.35 fF, DKO: 4.14 ± 0.44 fF, p<0.001). (E) Averaged FRET (Venus/mCerulean3) ratio time course from entire cell footprints, statistical analysis of baseline FRET ratio (F) (WT: 1.0315 ± 0.0065, DKO: 1.0228 ± 0.0067, p=0.38) and of FRET ratio change amplitude (G) (WT: 0.00487 ± 0.00075, DKO: −0.00031 ± 0.00065, p<0.001). Statistical analysis was performed using t-Test

## Discussion

### Pulse induced FRET dynamics resembles that of FRET changes preceding individual fusion events and is not consistent with diffusional equilibration

In our previous studies, a SCORE FRET ratio increase was specifically localized at sites of fusion events and preceded fusion by ∼90 ms in bovine chromaffin cells (8). The delay between the FRET increase and the fusion events indicates that the FRET increase reports the incorporation of SCORE into the SNARE complex in the vesicle priming step (7). In the current study cells were stimulated with voltage pulses, which depolarize the entire plasma membrane, inducing priming events at unknown locations, which may or may not be followed by fusion. When the 25 ms depolarization pulse elicits a calcium current (Fig. 2Ai), the Ca^2+^ influx induces a transient FRET ratio change indicating vesicle priming (Fig. 2C,E,G) as well as a membrane capacitance change indicating exocytotic fusion of a few vesicles (Fig. 2A,Aii). Priming events and fusion events in response to pulse stimulation may occur anywhere in the cell membrane. In our experimental approach, however, we performed TIR-FRET imaging in a large area of the cell footprint near the center of the cell and the detected priming events are therefore located in the cell’s footprint.

It is interesting to compare the SCORE FRET ratio change induced by 25 ms voltage pulses to the SCORE FRET ratio changes that were previously shown to precede individual fusion events recorded amperometrically using Electrochemical Detector Arrays (ECDs) (Fig. 4A) (8). It was found that the average SCORE FRET ratio increase in the central 4 pixel area surrounding the fusion site was followed by a plateau with a duration *t*_*Plateau*_=1.83 ± 0.11 s and a subsequent exponential decay with a time constant *τ*_*decay*_=1.56 ± 0.16s (Fig. 4B). The kinetics of the transient SCORE FRET ratio increase specifically localized at fusion sites and preceding fusion is thus very similar to that reported here for the change in the cell footprint in response to pulse stimulation of Ca^2+^ entry (*t*_*Plateau*_=2.6 ± 0.1 s, *τ*_*decay*_=1.56 ± 0.15 s).

**Fig. 4.**
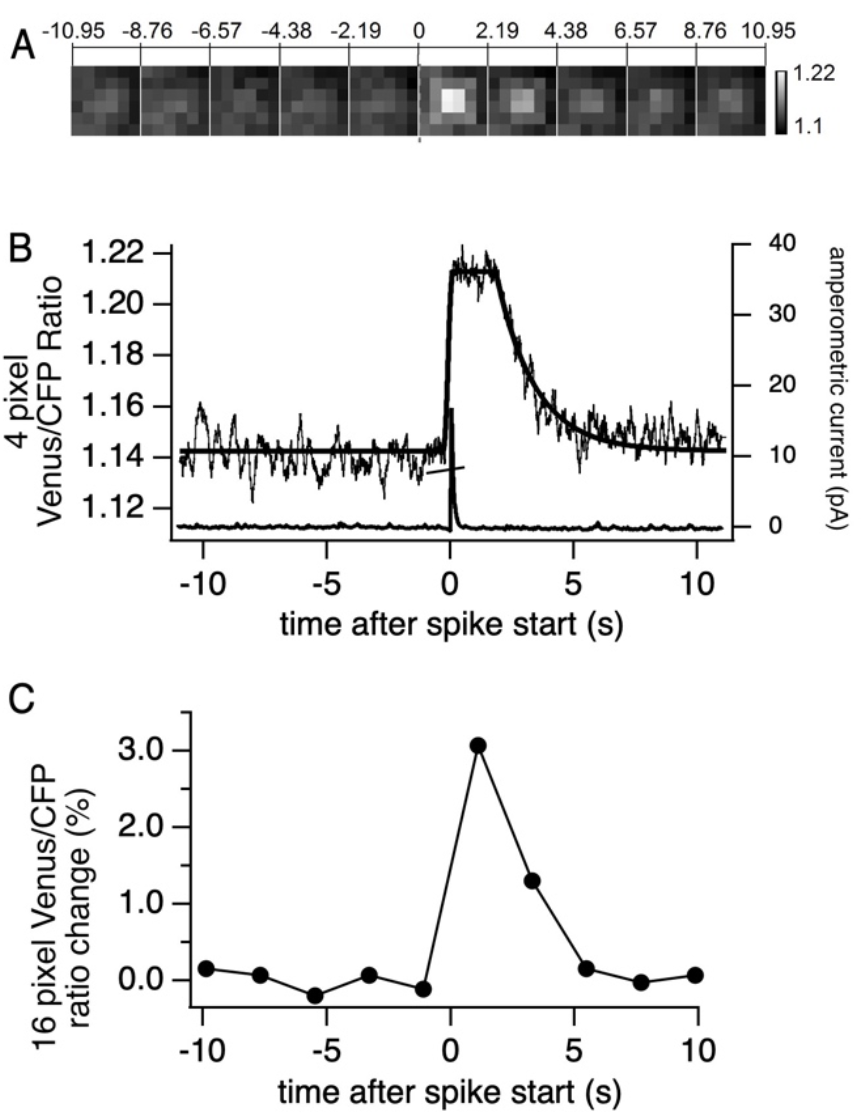
FRET dynamics correlated to individual fusion events. (A) Averaged FRET distributions in 6×6 pixel areas surrounding fusion sites from (8). (B) Averaged time course of FRET ratio change in central 2×2 pixel area aligned at amperometric spike onsets (black noisy trace from (8)). The averaged amperometric spike is shown as the grey trace. For comparison with the time course analysis of Fig. 2E, this data was now fitted (smooth curve, see text) with the same function as the data of Fig. 2E. (C) FRET ratio averaged over the central 4×4 pixels re-analyzed from the frames shown in panel A to account for the larger spread of the FRET ratio change, which exceeds the central 4×4 pixels as seen in (A).

In the previous studies using simultaneous SCORE TIR FRET imaging and amperometric ECD recordings (as in Fig. 4B), it remained unclear if the FRET ratio decrease was due to a reversal of the incorporation of SCORE in the SNARE complexes, i.e. SNARE complex disassembly, or if the assembled Hi-FRET SNARE complexes disperse after fusion, exchanging with unfolded Low-FRET SCORE.

The diffusion of Stx1 and SNAP25 in the plasma membrane has been characterized with high precision in an outstanding study using single molecule tracking in PC12 cells (19). The experiments showed that the majority of Stx1 and SNAP25 molecules undergo rapid diffusion with diffusion coefficients of ∼0.106 µm^2^/s and ∼0.25 µm^2^/s, respectively. However, about 1/3 of Stx1 molecules is sequestered in nanodomains and show much slower diffusion with a diffusion coefficient of ∼0.014 µm^2^/s. One in eight SNAP-25 molecules showed similarly slow diffusion with a diffusion coefficient of ∼0.015 µm^2^/s, indicating that these were bound to Stx1 copies with hindered diffusion (19).

In our previous experiments investigating SCORE FRET changes specifically related to fusion events, the SCORE or SCORE2 FRET changes extended over an area of 4×4 pixels (∼0.41 µm^2^) with most of the signal in the central 2×2 pixel area (7, 8) (see also Fig. 4). The time course of the FRET change was analyzed using the central 4 pixels with an area of ∼0.1 µm^2^. To estimate the expected fluorescence decay kinetics based on this area, we approximated the monitored area by a somewhat larger circle with radius R_C_=0.25 µm (area 0.2 µm^2^).

For a simplified estimate we assume that the SCORE copies changing to the high FRET state are generated at the center of a circular area with a radius of R_C_, we can calculate the probability of these copies in this circular area as a function of time. Based on 2-dimensional diffusion theory, the probability density of a molecule initially present at the center to move to a distance *r* at time *t* is

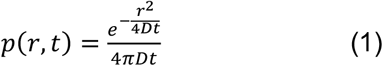

where *D* is the diffusion coefficient. The detail *p*(*r, t*) was plotted in the left panel of Fig. S1A. The probability *P*_*Circle*_ of the molecule to be present within a circle with radius *R*_*c*_ is then

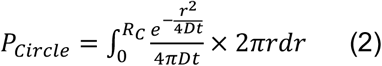

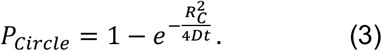

Thus, the half decay time t_½_ is obtained for *P*_*Circle*_ = 0.5 as

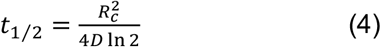

which is also plotted in the right panel of Fig. S1A.

The transition to the high FRET state is presumably associated with Stx1-SCORE complex formation or based on the result of Fig. 3, incorporation of SCORE into a ternary Stx1-SCORE-Syb2 complex. The predicted time course for a monitored circle with radius R_C_=0.25 µm (area 0.2 µm^2^) with the slow SNAP25 diffusion coefficient D=0.015 µm^2^ based on equation 3 and Eq. S14 is not exponential with a half decay time t_½_ ∼1.5 s (Fig. S1A). In a more precise estimate (SI Advanced Diffusion Estimates) we calculate the expected diffusional decay time based on the assumption that the high FRET copies are initially generated homogeneously inside the circle with radius R_C_=0.25 µm, which provides a slightly shorter t_½_=0.96 s (Eq. S20, Fig. S1B). This difference results from a larger fraction of SCORE copies that diffuse out of the circle region when they are initially generated homogenously within the circle rather than the center of the circle. Based on these estimates, the experimentally observed decay time constant of 1.56 s could thus be consistent with diffusional equilibration.

In the experiments with pulse stimulation monitoring the 30 µm^2^ area (Fig. 2) we approximate this area by a circle with radius R_C_=3 µm radius and the resulting time course of molecules leaving this area based on equation 3 is shown in Fig. S1A. This curve decays with a half time of 216 s (equation 3). Since for the larger area priming events presumably are randomly distributed, assuming initially homogeneous distribution (SI Advanced Diffusion Estimates) is presumably more appropriate (Fig. S1B) and provides t_½_ = 138 s. These estimates are about hundred times slower than the experimentally observed exponential decay with a time constant of 1.56 s (Fig. 2E).

Assuming the SNAP25 fast diffusion coefficient D = 0.24 µm^2^/s, the decay half time with the homogeneous generation assumption is ∼ 60 ms for R_C_ = 0.25 µm, and 8.6 s for R_C_ = 3 µm. These estimates show that the time course of the reversal of the SCORE FRET increase cannot be explained by diffusion, in particular since the experimentally measured FRET ratio decay is exponential and has the same time constant of ∼1.5 s for the small areas monitored during recordings at fusion sites and for the large footprint areas measured here with pulse stimulation. The results therefore clearly show that the reversal of the FRET change is not a consequence of SNARE complex dispersal after fusion. The FRET ratio decrease therefore reflects SCORE unfolding indicating SNARE complex disassembly. The disassembly begins at the end of the plateau, ∼1-3 s after assembly (Figs. 2E and 4B) see also (7, 8) and proceeds with a time constant of ∼1.5 s. The only caveat may be that after priming or fusion, ternary or tSNARE complexes may be endocytosed or otherwise translocated into the cell perpendicular to the cell membrane. We are, however, not aware of any studies indicating that SNAP25 or SNARE complexes are taken up into the cell following priming or fusion

### Some but not all pulse induced priming events are followed by fusion

To determine the average number of priming events in the cell footprint area, the FRET ratio changes in the cell footprint was compared to the FRET ratio changes of individual priming events preceding the corresponding fusion event. The average FRET ratio increase in the central 4 pixels surrounding fusion sites (Fig. 4A, B) is ∼6%. The increase in FRET ratio extends, however, over a larger 4×4 pixel (∼0.4 µm^2^) area (Fig. 3A). The average FRET ratio increase in this larger 16 pixel area is thus smaller, about 3.1% (Fig. 4C). To compare the FRET changes associated with individual events to the FRET ratio changes in response to pulse stimulation (Fig. 2) that were averaged over cell footprint areas of ∼1200 pixels (∼30 µm^2^), we need to quantify the FRET ratio change in the larger 4×4 pixel area that contributes to the FRET ratio change in the cell footprint. The expected FRET ratio change in this area from a 3.1% change in a 16 pixel (∼0.4 µm^2^) area surrounding a single fusion event is expected to be ∼0.04%. The average 0.2% change in FRET ratio in response to pulse stimulation would thus correspond to priming of ∼5 vesicles in the imaged footprint area. The average capacitance of a bovine chromaffin cell granule is 2.1 fF (20). If each priming event were followed by fusion, we would expect a capacitance change of ∼10 fF. The mean capacitance increase following a 25 ms depolarization pulse was ∼5-7 fF (Fig. 2Aii), corresponding to ∼3 chromaffin granules. It is, however, likely that additional priming events with the same SCORE structural change also occur elsewhere in the plasma membrane where the FRET ratio change is not recorded. The total number of vesicle priming events is therefore underestimated. Even if bovine chromaffin cells in culture may exhibit a strong polarization to release catecholamines towards their bottom (21) where the TIR FRET imaging also detects the priming events, the comparison indicates that only a fraction of all priming events induced by the 25 ms Ca^2+^ currents are followed by fusion.

### The priming step indicated by the SCORE FRET change depends on the presence of the vSNARE

Priming is a reversible dynamic process (22). Electron tomography studies suggest that vesicles dynamically transition between a loosely tethered state and a tightly docked state and that it is the tightly docked state from which vesicles can fuse and release transmitter (12, 13, 23). Tn chromaffin cells, this readily releasable pool (RRP) is regulated by intracellular Ca^2+^ ([Ca^2+^]_i_) In chromaffin cells the RRP has a maximal size around [Ca^2+^]_i_ =500 nM. At lower [Ca^2+^]_i_ priming decreases due to a decreased priming rate, at higher [Ca^2+^]_i_ priming decreases due to increased fusion rate, beginning to deplete the RRP (24). It has been suggested that after transient formation of a short four-helix bundle formed by the Stx1 and Syb2 SNARE domains and the helical segments Hd and He from the Stx1 linker, this small four-helix bundle opens up again to allow for SNAP25 to enter and complete the SNARE complex assembly (25). The incorporation of SNAP25 therefore appears to be the final step and it is this step, which defines the formation of the primed state (7). This model predicts that Syb2 must be bound to Stx1 when SNAP25 is incorporated to form the primed state and priming therefore requires the presence of the vSNARE. In support of this model our experiments using Syb2/Ceb DKO mice show directly that the SCORE FRET change induced by Ca^2+^ currents disappears when the vSNAREs are deleted. The result demonstrates directly that vesicle priming requires the presence of the vSNARE.

### Conclusion

The results presented here indicate that pulse stimulation of Ca^2+^ currents stimulates incorporation of SNAP25 into the SNARE complex, generating the primed state. Consistent with previous studies, this priming step requires the presence of vSNAREs. The primed state has a lifetime of a few seconds after which it is disassembled. This is the first characterization of the dynamics of priming and de-priming following Ca^2+^ current stimulation in live chromaffin cells.

## Materials and Methods

### Bovine chromaffin cell experiments

Bovine chromaffin cells were prepared as described (26) and SCORE expressed using the pSFV1 (Semliki Forest, Invitrogen) method (8). Cells expressing SCORE were voltage clamped in the whole cell patch clamp configuration while membrane capacitance was recorded using the Lindau-Neher technique (16). The standard extracellular solution contained (in mM): 140 mM NaCl, 5 mM KCl, 5 mM CaCl_2_, 1 mM MgCl_2_, 10 mM HEPES/NaOH, 20 mM glucose (pH 7.3). The pipette solution contained (in mM): 145 Cesium glutamate, 8 NaCl, 0.18 CaCl_2_, 0.28 BAPTA, 1 MgCl_2_, 2 ATP-Mg, 0.5 GTP-Na_2_, 0.3 cAMP, 10 HEPES-CsOH (pH 7.3), and 300 nM calculated free [Ca^2+^]_i_. A Zeiss Axiovert 135 TV microscope modified for objective-type TIRF illumination as described (8, 27) and equipped with a Zeiss Plan-Fluar 1.45 NA 100X oil objective was used for simultaneous TIRF imaging. Illumination was provided by a mercury arc lamp (X-cite, 120PC) through a 436/10nm excitation filter and a 455 dichroic. Fluorescence emission from an ∼30 µm^2^ area of the footprint of the cell was imaged simultaneously in the CFP (465-495 nm) and Venus (520-550 nm) bands with an image splitter (Dual View, Optical Insights) using a 505 nm dichroic.

Cells were stimulated with 25 ms pulses from −70 mV to +10 mV (not corrected for liquid junction potentials) to activate calcium currents. Since we did not use Na^+^ channel blockers, the first 5 ms after pulse start were dominated by Na^+^ current. For quantification of the Ca^2+^ current, only the current from 5 to 25 ms was therefore averaged and the initial 5 ms were excluded. For each recording, image sequences (2,000 frames) were collected with 100 ms exposure time and 1.7928 ms readout interval using an EMCCD camera (Andor iXon) and its accompanying software such that one frame was acquired every 101.7928 ms simultaneously for both channels. Stimulation pulses were applied at random times during the recording at ∼15 s interpulse intervals. The camera provides a TTL signal indicating exposure times that was recorded to synchronize the timing of the fluorescence image frames relative to the stimulation pulses for analysis. Fluorescence intensities were obtained in camera units after background subtraction. The background fluorescence intensity was measured in a region outside the cell. The ratio of background subtracted areas of the cell footprint in the Venus and CFP channels was determined, generating a frame-by-frame FRET ratio record.

### Kinetic analysis of averaged FRET time course induced by pulse depolarization

The time resolution of the amperometric spikes reported by the ECD events is 1ms. Due to low signal-to-noise ratio of the fluorescence images, however, imaging frame durations of ∼100 ms are needed to collect sufficient signal photons such that the precise temporal correlation of the FRET change with the time of fusion pore formation is uncertain. To obtain a more precise time correlation we used the Event COrrelation Microscopy (ECOM) method to analyze imaging signals with sub-frame time super-resolution (8, 17). In brief, we padded the frame intensities with 1-ms time points, assigning the same frame intensity to all the points within that frame. This makes it possible to align the imaging data at the onset of the amperometric ECD spikes with 1 ms precision such that the frames from different recordings can be temporally aligned with the respective amperometric spikes with sub-frame resolution including a time interval from 10s before pulse start to 10s after pulse start. The temporally aligned padded traces were averaged providing the averaged FRET time course, representing the average time course of the FRET change lowpass filtered with the ECOM step response function for the specific exposure and read-out times used in the recordings (17). These averaged traces were fitted with the ECOM step response function (Equation 9 in ref (17)), which was modified by adding an exponential decrease back to baseline to account for the decrease following the plateau. In addition to the amplitude and time of the FRET ratio increase, this fit thus provided also the time for the end of the plateau, *t*_*plateau*_, where the exponential decay starts and the time constant of the exponential decay *τ*_*decay*_.

### Syb2/Ceb DKO mouse embryonal chromaffin cell experiments

Syb2/Ceb DKO or wild type littermate mouse E17-E18 embryos were obtained and chromaffin cells isolated and plated as described (18, 28). Recombinant Semliki Forest Virus was used to introduce SCORE2 into the cells after 2 days in culture (7). Cell membrane capacitance was recorded in the patch clamp whole-cell configuration using a Patchmaster controlled EPC-10 amplifier (HEKA electronics).. Cells were stimulated with 200 ms pulses from −70 mV to +10 mV (not corrected for liquid junction potentials). The depolarization evoked exocytosis was detected as change in cell capacitance, estimated by the Lindau-Neher technique (16). Membrane capacitance (Cm) changes were analyzed with customized IgorPro routines (WaveMetrics). The standard extracellular solution contained (in mM): 140 mM NaCl, 5 mM KCl, 5 mM CaCl_2_, 1 mM MgCl_2_, 10 mM HEPES/NaOH, 20 mM glucose (pH 7.3). The pipette solution contained (in mM): 145 Cesium glutamate, 8 NaCl, 0.18 CaCl_2_, 0.28 BAPTA, 1 MgCl_2_, 2 ATP-Mg, 0.5 GTP-Na_2_, 0.3 cAMP, 10 HEPES-CsOH (pH 7.3), and 300 nM calculated free [Ca^2+^]_i_.

Fluorescence imaging was performed in TIRF Mode as described (7) with an inverted microscope Olympus IX83 equipped with a 1.45 NA 100x objective and an Optosplit III (Cairn Research) image splitter fitted with a 509nm dichroic and emission filters at 467-499 nm for the mCerulean3 and 520-550nm for the Venus channel. The images of the two channels were projected side-by-side onto an EMCCD (Andor iXon3) run by its accompanying software (Solis) with 100 ms exposure time starting 10 s before stimulation and ending 10 s after stimulation. Illumination in TIRF mode was provided by a 442nm laser (Changchun New Industries Optoelectronics Tech.Co.).

## Acknowledgements

This work has been supported by US NIH grants R01GM121787 and R35GM139608 and European Research Council grant ADG 322699.

## Author contributions

M.L designed the project, Q.F, Y.Z and M.L performed the experiment and analyzed the data, D.A. performed the numerical diffusion calculations and all authors wrote the manuscript.

## Declaration of generative AI and AI-assisted technologies in the writing process

During the preparation of this work the third author used ChatGPT in order to improve language and readability. After using this tool/service, the author(s) reviewed and edited the content as needed and took full responsibility for the content of the publication.

