## Supplemental information for "The dynamics of SNARE complex assembly and disassembly in response to Ca^2+^ current activation in live chromaffin cells"

### Numerical simulations to calculate the fraction of SCORE copies retained in the circle region

Given that the high FRET state of SCORE copies are generated at $t = 0$ at the center of a circular area with a radius $R_C$ initially stayed inside the circle region. Then, it is freely diffused 2 dimensionally with a diffusion coefficient of $D$.

The governing equation that describes the probability density function of the molecule with respect to the radius coordinate $r$ and time $t$ is:

$$\frac{\partial p}{\partial t} = \frac{D}{r}\frac{\partial}{\partial r}\left(r\frac{\partial p}{\partial r}\right) \ldots\ldots\ldots\ldots\ldots\ldots\ldots\ldots\ldots\ldots\ldots\ldots\ldots. \text{Eq. S1.}$$

The boundary conditions are:

$$p(r \to +\infty, t) = 0, \ldots\ldots\ldots\ldots\ldots\ldots\ldots\ldots\ldots\ldots\ldots\ldots\ldots. \text{Eq. S2}$$

$$\text{and } \left.\frac{\partial p}{\partial r}\right|_{r=0} = 0, . \ldots\ldots\ldots\ldots\ldots\ldots\ldots\ldots\ldots\ldots\ldots\ldots\ldots \text{Eq. S3.}$$

These equations mean that the probability that high FRET state of SCORE copies appear far away from the center of the circle region is 0 (Eq. S2) and the cylindrical symmetry (Eq. S3).

By assuming that the high FRET state of SCORE copies generated at the center of the region, the initial condition is

$$p(r, 0) = \begin{cases} +\infty \ (r = 0) \\ 0 \quad (r > 0) \end{cases}, \ldots\ldots\ldots\ldots\ldots\ldots\ldots\ldots\ldots\ldots\ldots.. \text{Eq. S4,}$$

whereas the assumption that the high FRET state of SCORE copies generated homogeneously in the circle region is

$$p(r, 0) = \begin{cases} 1/\pi R_c^2 \ (r < R_c) \\ \quad 0 \quad (r \geq R_c) \end{cases}, \ldots\ldots\ldots\ldots\ldots\ldots\ldots\ldots\ldots\ldots\ldots.. \text{Eq. S5.}$$

The probability density $p(r, t)$ always satisfies

$$\int_0^{+\infty} 2\pi p(r, t) r dr = 1. . \ldots\ldots\ldots\ldots\ldots\ldots\ldots\ldots\ldots\ldots\ldots.. \text{Eq. S6.}$$

### Scaling variables to build the dimensionless model

Then, we scaled Eq. S1 by applying

$$\bar{r} = \frac{r}{R_c},$$

which is the dimensionless radius coordinate scaled by the radius of the circular region. We get

$$\frac{\partial p}{\partial t} = \frac{\mathrm{D}}{R_c^2}\frac{1}{\bar{r}}\frac{\partial}{\partial \bar{r}}\left(\bar{r}\frac{\partial p}{\partial \bar{r}}\right). \dots\dots\dots\dots\dots\dots\dots\dots\dots\dots\dots\dots \text{Eq. S7.1.}$$

Then to reduce the time dimension, we apply

$$\tau = \frac{t}{t_\mathrm{c}} \text{ where } t_\mathrm{c} = \frac{R_c^2}{D},$$

We get

$$\frac{1}{t_c}\frac{\partial p}{\partial t} = \frac{1}{t_c}\frac{1}{\bar{r}}\frac{\partial}{\partial \bar{r}}\left(\bar{r}\frac{\partial p}{\partial \bar{r}}\right). \dots\dots\dots\dots\dots\dots\dots\dots\dots\dots\dots\dots \text{Eq. S7.2}$$

to eliminate the dimensions in the model. Eq. S7.2 can be simplified by cancelling the $1/t_0$ term at both hand sites and results in

$$\frac{\partial p}{\partial \tau} = \frac{1}{\bar{r}}\frac{\partial}{\partial \bar{r}}\left(\bar{r}\frac{\partial p}{\partial \bar{r}}\right). \dots\dots\dots\dots\dots\dots\dots\dots\dots\dots\dots\dots\dots. \text{Eq. S7.3.}$$

The boundary conditions become

$$p(\bar{r} \to +\infty, \tau) = 0, \dots\dots\dots\dots\dots\dots\dots\dots\dots\dots\dots\dots. \text{Eq. S8}$$

$$\text{and } \left.\frac{\partial p}{\partial \bar{r}}\right|_{\bar{r}=0} = 0, \dots\dots\dots\dots\dots\dots\dots\dots\dots\dots\dots\dots\dots \text{Eq. S9.}$$

Thus, the two initial conditions in Eq. S5 and Eq. S6 become

$$p(\bar{r}, 0) = \begin{cases} +\infty \ (\bar{r} = 0) \\ 0 \quad (\bar{r} > 0) \end{cases}, \dots\dots\dots\dots\dots\dots\dots\dots\dots\dots\dots.. \text{Eq. S10}$$

$$\text{and } p(\bar{r}, 0) = \begin{cases} 1/\pi \ (\bar{r} < 1) \\ 0 \quad (\bar{r} \geq 1) \end{cases}., \dots\dots\dots\dots\dots\dots\dots\dots\dots\dots\dots. \text{Eq. S11.}$$

The probability density $p(\bar{r}, \tau)$ always satisfies

$$\int_0^{+\infty} 2\pi p(\bar{r}, \tau)\bar{r} d\bar{r} = 1.. \dots\dots\dots\dots\dots\dots\dots\dots\dots\dots\dots.. \text{Eq. S12.}$$

By scaling the model, we could find the general shape of diffusive profile.

### Solving the model numerically and compared with analytical results

By solving Eq. S7.3 with the initial condition S10, the probability density is

$$p(\bar{r},\tau) = \frac{e^{-\frac{\bar{r}^2}{4\tau}}}{4\pi\tau} \ldots \ldots \ldots \ldots \ldots \ldots \ldots \ldots \ldots \ldots \ldots \ldots \ldots \ldots \ldots \ldots \text{Eq. S13},$$

which is also shown in the left panel of Fig. S1A, and

$$P_{Circle}(\tau) = \int_0^1 2\pi p(\bar{r},\tau)\bar{r}d\bar{r} = \int_0^1 2\pi \times \frac{e^{-\frac{\bar{r}^2}{4\tau}}}{4\pi\tau}\bar{r}d\bar{r} = 1 - e^{-\frac{1}{4\tau}} \ldots \ldots \ldots \ldots \ldots \text{Eq. S14}.$$

shown in the right panel of Fig. S1A.

Thus,

$$t_{1/2} = \frac{t_c}{4\ln 2} \approx 0.36t_c = \frac{0.36R_C^2}{D} \ldots \ldots \ldots \ldots \ldots \ldots \ldots \ldots \ldots \ldots \ldots \text{Eq. S15}.$$

By solving Eq. S7 with the initial condition S11, the probability density is shown in the left panel of Fig. S1B.

By integrating the dimension $\bar{r}$ in Eq. S7 from 0 to 1 and multiply by $2\pi$, from the left-hand side, we get

$$2\pi \int_0^1 \frac{\partial p}{\partial \tau}\bar{r}d\bar{r} = \frac{d}{d\tau}\int_0^1 2\pi p(\bar{r},\tau)\bar{r}d\bar{r} = \frac{dP_{Circle}(\tau)}{d\tau} \ldots \ldots \ldots \ldots \ldots \ldots \ldots \ldots \text{Eq. S16},$$

whereas from the right-hand side, we get

$$\int_0^1 2\pi \frac{1}{\bar{r}}\frac{\partial}{\partial \bar{r}}\left(\bar{r}\frac{\partial p}{\partial \bar{r}}\right)\bar{r}d\bar{r} = 2\pi\left(\bar{r}\frac{\partial p}{\partial \bar{r}}\right)\Big|_0^1 = 2\pi\frac{\partial p}{\partial \bar{r}}\Big|_{\bar{r}=1} \ldots \ldots \ldots \ldots \ldots \text{Eq. S17}.$$

i.e.

$$\frac{dP_{Circle}(\tau)}{d\tau} = 2\pi\frac{\partial p}{\partial \bar{r}}\Big|_{\bar{r}=1} \ldots \ldots \ldots \ldots \ldots \ldots \ldots \ldots \ldots \ldots \ldots \ldots \ldots \text{Eq. S18}.$$

So $\frac{dP_{Circle}(\tau)}{d\tau}$ is proportional to the probability density gradient at the boundary of the region.

Thus,

$$P_{\text{inside}}(\tau) = \int_0^\tau \frac{dP_{Circle}}{d\tau'}d\tau' = \int_0^\tau 2\pi\frac{\partial p}{\partial \bar{r}}\Big|_{\bar{r}=1}(\tau')d\tau' \ldots \ldots \ldots \ldots \ldots \ldots \ldots \ldots \text{Eq. S19}.$$

which is the semi-analytical solution of $P_{\text{inside}}(\tau)$ shown in the right panel of Fig. S1B.

This gives the half decay time is

$$t_{1/2} \approx 0.23 t_c = \frac{0.36 R_C^2}{D} \ldots\ldots\ldots\ldots\ldots\ldots\ldots\ldots\ldots\ldots\ldots\ldots\ldots \text{Eq. S20.}$$

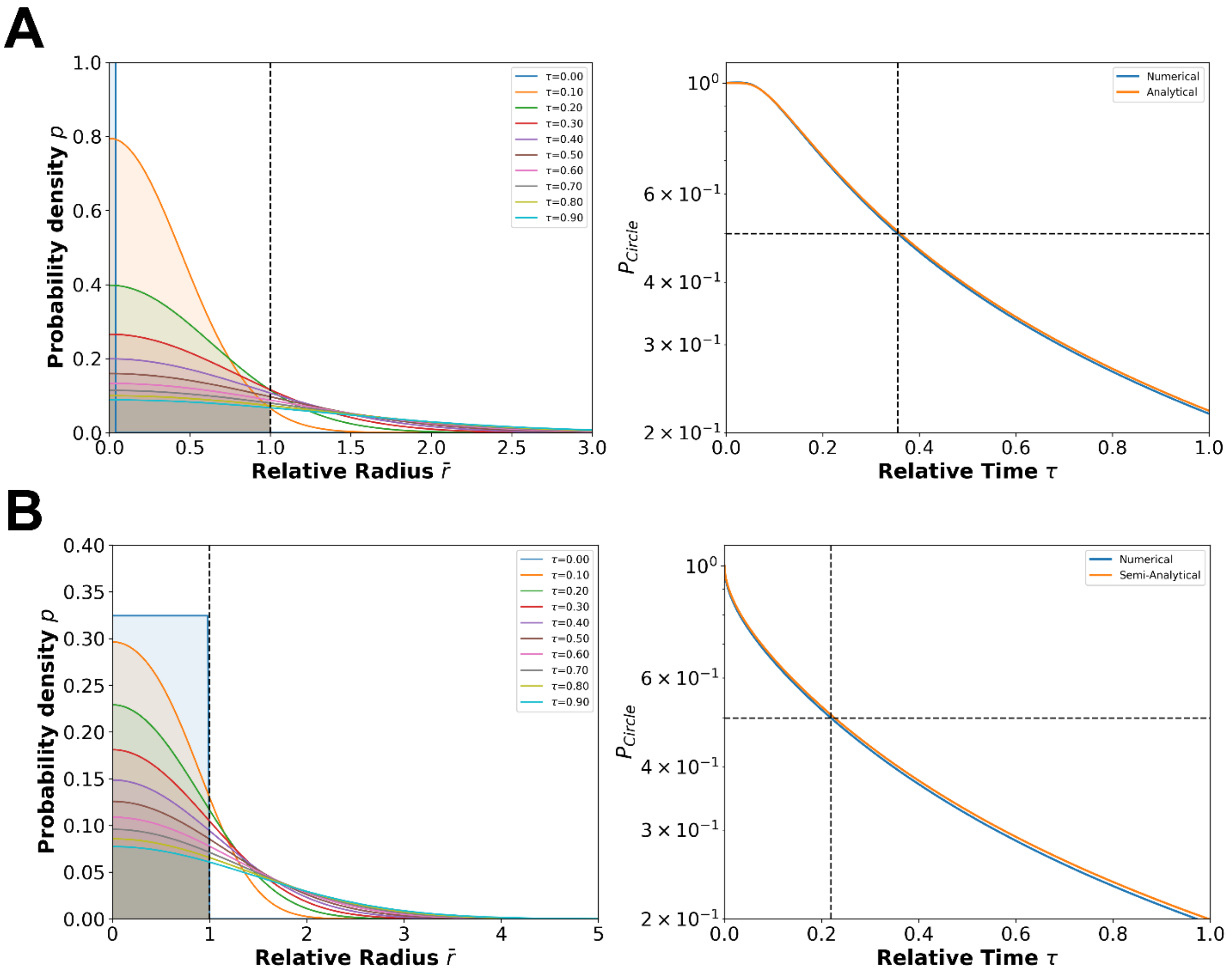


Figure S1. Numerical solution of the probability density for the high-FRET state of SCORE copies and the probability that a high-FRET state SCORE copy remains inside the circular region.

(A) (Left) Probability density profile $p(\bar{r}, \tau)$ as a function of the relative radius coordinate $\bar{r} = r/R_C$ at relative time $\tau = t/t_x$ where $t_c = R_c^2/D$, calculated using the point-source initial condition in Eq. S10. The solution, with $\bar{r}$ ranging from 0 to 20 and $\tau$ from 0 to 1, was obtained using the finite difference method, discretizing the region with a dimensionless radial step $\delta\bar{r} = 0.02$ and a dimensionless time step $\delta\tau = 0.0001$. The vertical dashed line indicates the relative radius coordinate of the circular region boundary. The areas of the shaded regions are the probability of high-FRET state of

SCORE copies remain in the circle region at different relative time $\tau$. Note: the profile $p(\bar{r}, 0)$ shown in this panel is truncated since the peak value is too high to fit the scale of the panel. (Right) The probability $P_{Circle}$ was computed numerically as the areas of the shaded regions in the left panels using Eq. S14 and shows excellent agreement with the analytical solution derived from Eq. S14. The result also yields the expression for $t_{1/2}$ given in Eq. 15. Dash horizonal line: $P_{Circle} = 0.5$, and vertical line: $\tau_{1/2} \approx 0.36$, which gives $t_{1/2} \approx 0.36 R_C^2/D$. (B) (Left) Probability density profile $p(\bar{r}, \tau)$ as a function of the relative radius coordinate $\bar{r}$ at relative time $\tau$, calculated using the homogeneous dispersion initial condition in Eq. S11. The solution, with $\bar{r}$ from 0 to 20 and $\tau$ from 0 to 1, was also obtained using the finite difference method with the same $\delta\bar{r}$ and $\delta\tau$ as in (A). The vertical dashed line indicates the relative radius coordinate of the circular region boundary. The areas of the shaded regions are the probability of high-FRET state of SCORE copies remain in the circle region at different relative time $\tau$. (Right) The probability $P_{Circle}$ was computed both numerically using Eq. S14 and semi-analytically using Eq. S19 with the numerical concentration profiles. It demonstrates strong agreement between the numerical and semi-analytical solutions, ultimately yielding the expression for $t_{1/2}$ in Eq. 20. Dash horizonal line: $P_{Circle} = 0.5$, and vertical line: $\tau_{1/2} \approx 0.23$, which gives $t_{1/2} \approx 0.23 R_C^2/D$.
